# The ER chaperones BiP and Grp94 regulate the formation of insulin-like growth factor 2 (IGF2) oligomers

**DOI:** 10.1101/2020.09.24.311779

**Authors:** Yi Jin, Judy L.M. Kotler, Shiyu Wang, Bin Huang, Jackson C. Halpin, Timothy O. Street

## Abstract

While cytosolic Hsp70 and Hsp90 chaperones have been extensively studied, less is known about how the ER Hsp70 and Hsp90 paralogs (BiP and Grp94) recognize clients and influence their folding. Here, we examine how BiP and Grp94 influence the folding of insulin-like growth factor 2 (IGF2). Full-length proIGF2 is composed of an insulin-like hormone and an E-peptide that has sequence characteristics of an intrinsically disordered region. We find that the E-peptide region allows proIGF2 to form oligomers. BiP and Grp94 influence both the folding and the oligomerization of proIGF2. BiP and Grp94 exert a similar holdase function on proIGF2 folding by preferentially binding the proIGF2 unfolded state, rather than stabilizing specific folding intermediates and changing the proIGF2 folding process. In contrast, BiP and Grp94 exert counteracting effects on proIGF2 oligomerization. BiP suppresses proIGF2 oligomerization under both ADP and ATP conditions. Interestingly, Grp94 can enhance proIGF2 oligomerization when Grp94 adopts an open conformation (ADP conditions), but not when Grp94 is in the closed conformation (ATP conditions). We propose that BiP and Grp94 regulate the assembly of proIGF2 oligomers, and that regulated oligomerization may enable proIGF2 to be effectively packaged for export from the ER to the Golgi.

## Introduction

Hsp70 and Hsp90 chaperones maintain protein quality control in the cytosol, while the endoplasmic reticulum (ER) and mitochondria have their own homologous Hsp70/Hsp90 systems (ER: BiP/Grp94, mitochondria: Mortalin/Trap1). Hsp70 and Hsp90 family members require ATP binding and hydrolysis to cycle through distinct conformational states with distinct client binding properties^1-7^. Both chaperones are heavily regulated by co-chaperones and post-translational modifications^8-9^. Adding to this complexity, Hsp70 and Hsp90 often associate during the chaperoning process, which can make their functions interconnected^10-11^.

Client interactions with cytosolic Hsp90 have been extensively studied^6, 12-15^, however less is known about how Grp94 and Trap1 recognize clients and subsequently influence their folding and activity. For example, while cytosolic Hsp90 can target clients in monomeric and oligomeric states ^12-14, 16^, it is not clear whether Grp94 operates in a similar manner. Cytosolic Hsp90 has been proposed to be specialized for the late-stages of folding for some client proteins^17-19^, in contrast to Hsp70 that canonically acts as a holdase by binding fully unfolded clients. However, experimental tests of whether Hsp90 has preferential recognition of partially folded states has yielded inconsistent conclusions^20-22^, likely due to the technical difficulty in identifying client folding intermediates and quantifying their populations. It is unknown whether Grp94 acts as a holdase by stabilizing fully unfolded states or whether it can actively influence a folding process.

Here we examine interactions between Grp94, BiP, and IGF2. IGF2 is one of only small collection of ER proteins that are known to stringently depend on Grp94^23^. Indeed, Grp94 is involved in the successful secretion of all members of the IGF family (insulin, IGF1 and IGF2)^24-27^, but the underlying mechanism is not understood. Grp94 mutations that abolish binding of ATP or are defective in hydrolysis cannot support cell viability in serum-free media due to the loss of secreted IGF2^25-26^. In contrast to the limited clientele of Grp94, BiP exhibits broad specificity due to the ubiquity of the short hydrophobic motifs that it recognizes^28^.

IGF proteins have three disulfide bonds in their hormone region. Disulfide-bonded proteins are uniquely well-suited for studying folding pathways because their folding intermediates can be trapped by quenching techniques and separated by reverse-phase HPLC (RP-HPLC). These methods have been applied with success to robust folding systems, like bovine pancreatic trypsin inhibitor (BPTI)^29-30^, but little is known about whether the same set of techniques can reveal the folding properties of proteins that are in the process of receiving chaperone assistance.

Mouse IGF2 is composed of a 24-residue signal sequence followed by a 67-residue hormone region and an 87-residue E-peptide region^31^ (Figure 1). ProIGF2 (the hormone region and E-peptide region) does not have N-linked glycosylation sites^32^, thus it primarily depends on the non-lectin chaperone machinery. Upon folding in the ER, proIGF2 is packaged into vesicles for transport to the Golgi^33^. Upon entry into the Golgi, proIGF2 is proteolytically separated into mature IGF2 (mIGF2) and the E-peptide^34^. *In-vitro* folding studies of IGF1 & IGF2 have focused on the mature constructs^35-37^. Here we focus on full-length proIGF2 because it is the biologically-relevant construct for BiP and Grp94.

**Figure 1.**
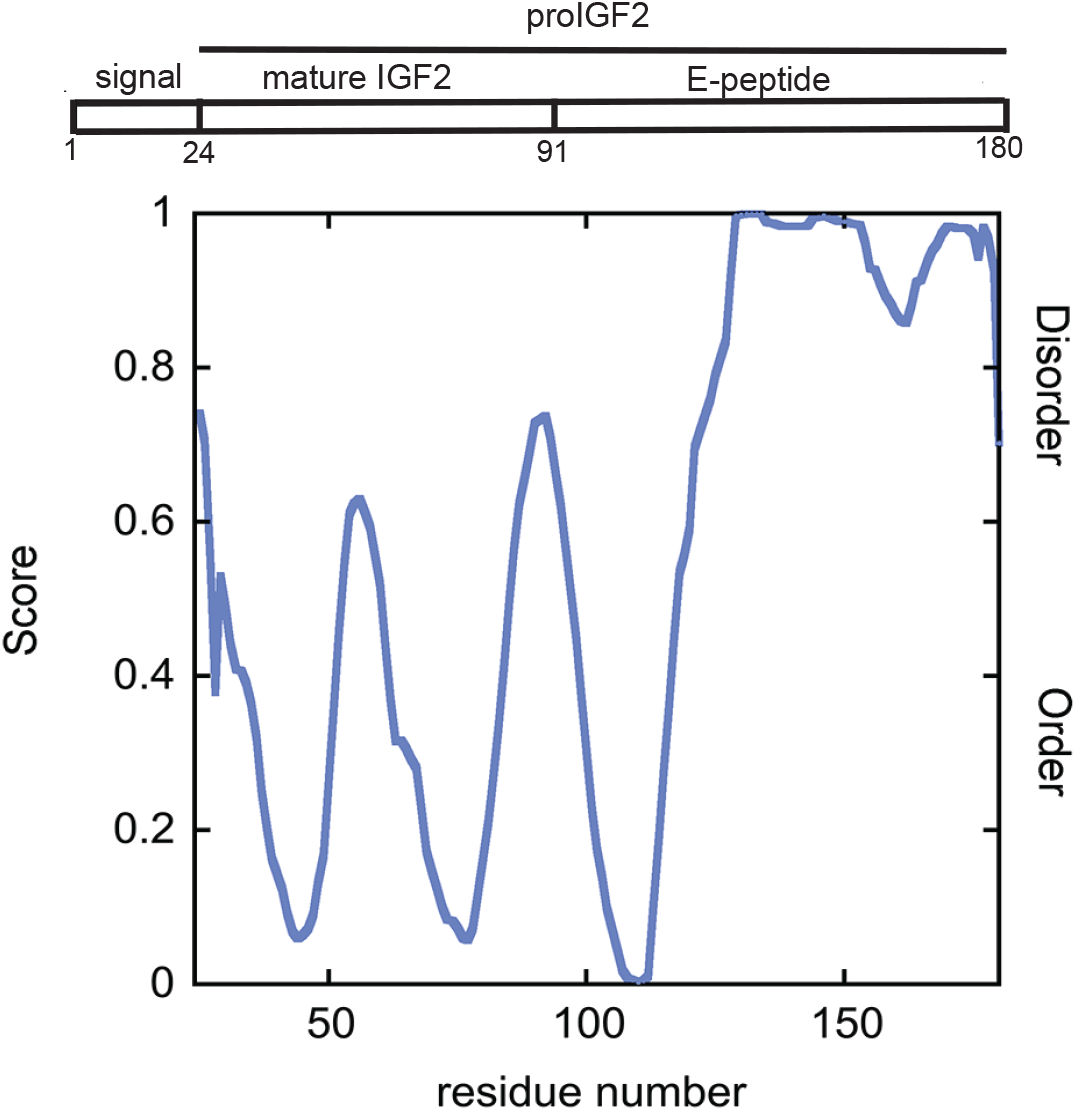
IGF2 E-peptide is an intrinsic disordered region. Sequence composition of IGF2 and disorder prediction of proIGF2 by PONDR-VLXT ^64^.

Little is known about the biological role of the E-peptide region, although it is predicted to be intrinsically disordered based on primary sequence (Figure 1). Intrinsically disordered proteins (IDPs) have a strong propensity to form condensates ^38-40^, which are dynamic soluble oligomers that can transition into irreversible aggregates. Chaperones are believed to be involved in maintaining the dynamics and solubility of condensates^41-42^, however this is an active area of research with many open questions.

Here we find that Grp94 is not specialized for the late-stages of proIGF2 folding. Rather, both BiP and Grp94 exert a holdase function by preferentially interacting with the unfolded state of proIGF2. However, we find that the E-peptide region promotes proIGF2 oligomerization, and that Grp94 and BiP exert counteracting effects on the formation of proIGF2 oligomers.

## Results

Under denaturing conditions proIGF2 can be fully reduced by TCEP (Supplemental Figure 1A) and subsequent experiments are initiated by diluting proIGF2 out of denaturant. After dilution proIGF2 maintains a fully reduced state under TCEP conditions, whereas proIGF2 folds under oxidizing conditions using a mixture of oxidized and reduced glutathione. Figure 2A shows how proIGF2 folding can be monitored by RP-HPLC (see Methods for details). Folding and disulfide formation are quenched by phosphoric acid. The quenched samples remain soluble and do not change their disulfide status for at least 20h (data not shown). A commercial standard of mIGF2 has a similar retention time as the major population of the recombinant refolded mIGF2 (Supplemental Figure 1B), indicating that proper refolding has occurred.

**Figure 2.**
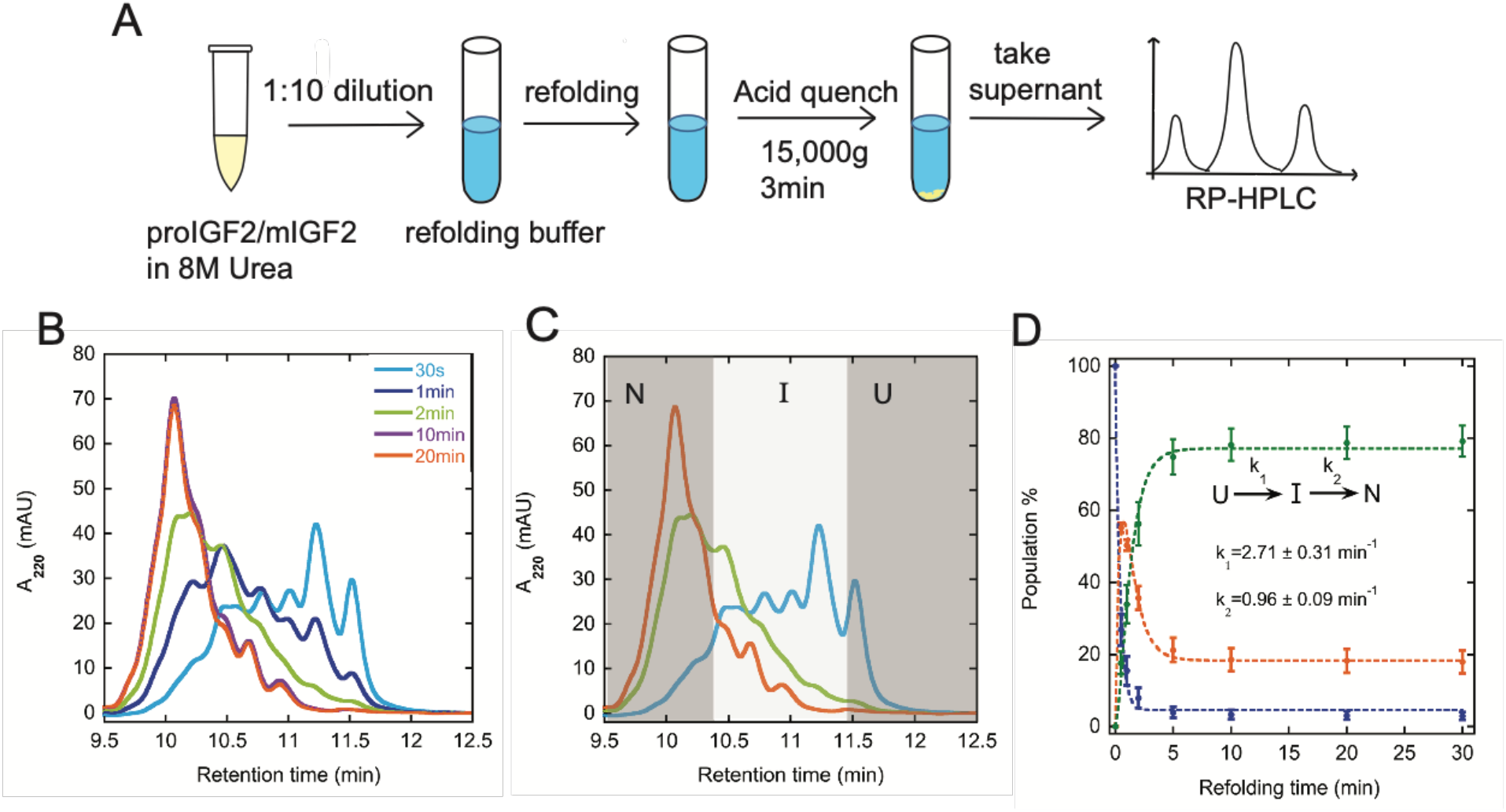
ProIGF2 is capable to fold. **A)** Protocol to populate and characterize proIGF2 folding states by RP-HPLC. **(B)** HPLC chromatograms of proIGF2 folding intermediates quenched at various folding times. Folding times shown in the legend. **(C)** ProIGF2 folding is categorized into three states (U, I, and N). **(D)** Populations of proIGF2 were fitted with a three-state model (see Methods). The populations of U, I and N are color-coded in blue, orange, and green respectively. Error bars are the S.E.M of at least three measurements. Refolding buffer: 50mM Tris pH 7.5, 100mM KCl, 1mM ATP, 1mM MgCl_2_, 1.25mM GSSG, 6.25 mM GSH, 0.8M urea, 37°C.

Because proIGF2 folding is redox-dependent, we expect strongly pH-dependent folding kinetics whereby disulfide formation should be accelerated at higher pH values, and proIGF2 indeed exhibits this expected pH-dependence (Supplemental Figure 1C). We also tested a range of redox potentials and observe faster folding kinetics and higher solubility under more oxidizing conditions (Supplemental Figure 1D). We selected pH 7.5 and a redox potential of 5.0 (ratio of GSH/GSSG) for more detailed analysis of proIGF2 folding. Under these conditions proIGF2 undergoes a modest level of misfolding and aggregation. For example, after 30 minutes of folding under these conditions the soluble fraction of proIGF2 is 72±7% of the starting material. Thus, these conditions allow for potential improvements in proIGF2 folding to be observed from the influence of BiP and Grp94.

Multiple folding states of proIGF2 and mIGF2 are populated during refolding (Figure 2B, Supplemental Figure 1B). While a complete folding analysis is not feasible due to overlapping chromatography peaks, a simplified folding description can be achieved by combining elution peaks into three groups based on their retention times (Figure 2C & Supplemental Figure 1E, see Methods for details). We find that proIGF2 and mIGF2 follow similar folding kinetics (Figure 2D & Supplemental Figure 1F), indicating that the E-peptide region does not have a strong influence on the folding of the mature region.

### BiP and Grp94 slow down proIGF2 folding by stabilizing the unfolded state

BiP and Grp94 could influence proIGF2 folding through a wide variety of mechanisms. For example, BiP and Grp94 could accelerate proIGF2 folding, selectively bind specific vulnerable folding intermediates, or even change the folding pathway to avoid vulnerable folding intermediates. The well-populated proIGF2 folding intermediates provide an opportunity to distinguish between these scenarios.

Both BiP and Grp94 decelerate proIGF2 folding (Figure 3). BiP and Grp94 stabilize the proIGF2 unfolded state (11.6 minutes retention time) as is evident by its increased population (see 0.5min folding time). The accumulation of proIGF2 folding intermediates is decelerated, for example the intermediate at 11.2 minutes retention time (see 1min refolding time). Finally, the build-up of fully-oxidized proIGF2 at 10.1 minutes retention time is also decelerated (see 2min refolding time). No new highly populated folding intermediates are observed. Both BiP and Grp94 have a minimal influence on proIGF2 refolding yield (Supplemental Figure 2).

**Figure 3.**
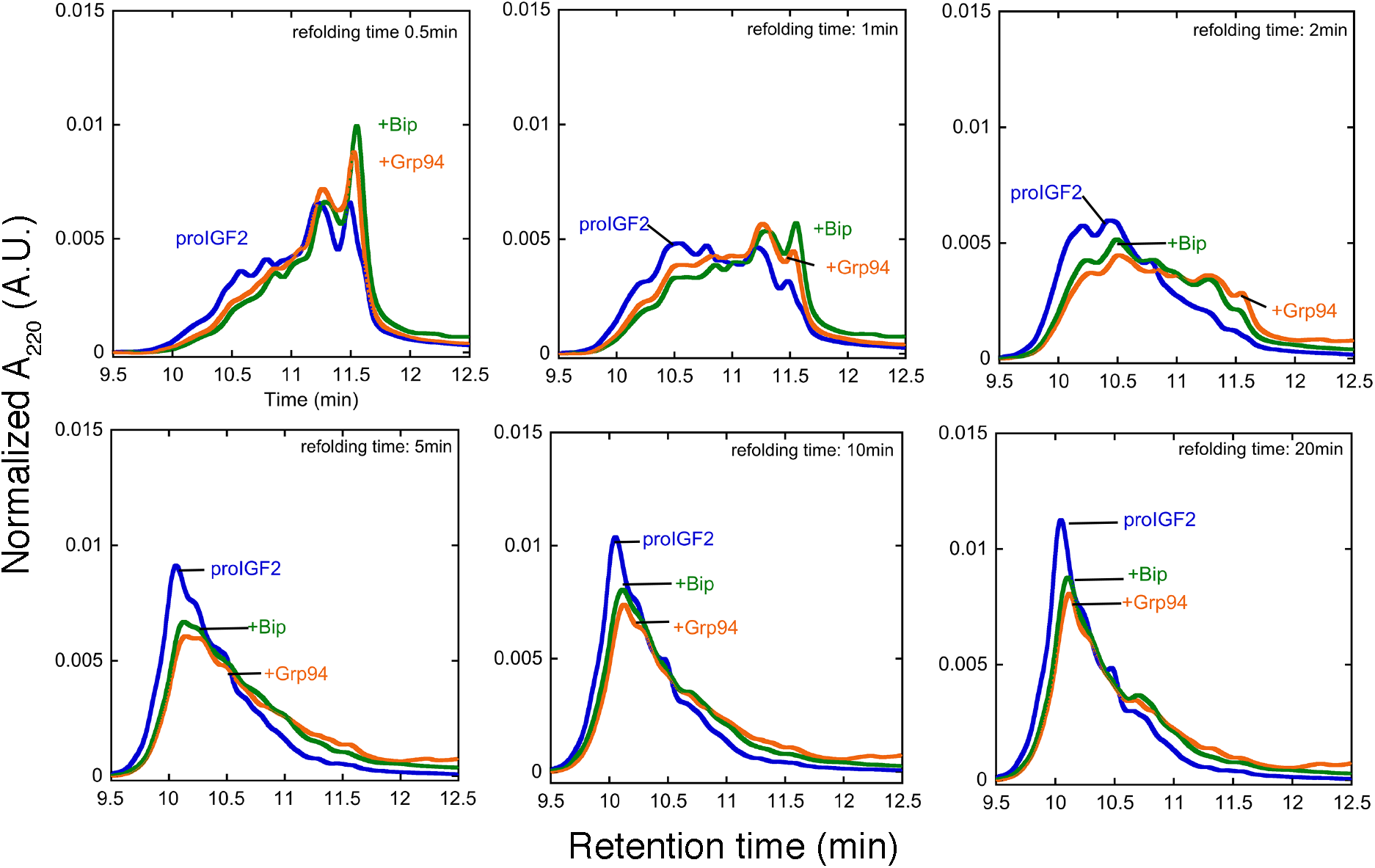
BiP and Grp94 bind to unfolded proIGF2. Influence of BiP and Grp94 on the folding of proIGF2. HPLC elution profiles of proIGF2 with and without chaperone are compared at various refolding times. Peak areas are normalized for a direct comparison between different conditions. Refolding buffer conditions are the same as Figure 2.

The above results are qualitatively consistent with BiP and Grp94 decelerating folding by binding to unfolded proIGF2 and preventing disulfide formation in the bound state. To test this idea quantitatively, we determined the build-up rate of folding intermediates and their progression to a fully oxidized state (Supplemental Figure 3) as was done for proIGF2 alone in Figures 2C&D. The kinetic data were fit with three models: 1) BiP and Grp94 bind to the proIGF2 unfolded state; 2) BiP and Grp94 bind proIGF2 folding intermediates; 3) BiP and Grp94 bind to the proIGF2 unfolded state and folding intermediates (see Appendix for details). Model 1 provides a better fit than Model 2. Model 3 has more fitting parameters but provides only a slight improvement over Model 1. We conclude that the influence of BiP and Grp94 on proIGF2 refolding kinetics is simply due to preferential binding to unfolded proIGF2.

While the above results show BiP and Grp94 exerting a simple holdase function on the folding of proIGF2, we also observe that BiP and Grp94 ATPase activity are both increased by proIGF2 (Supplemental Figure 4), which suggests that these chaperones could be actively influencing some other aspect of proIGF2 behavior. Indeed, as described next proIGF2 can oligomerize due to the influence of the E-peptide, and that BiP and Grp94 influence the oligomerization process.

**Figure 4.**
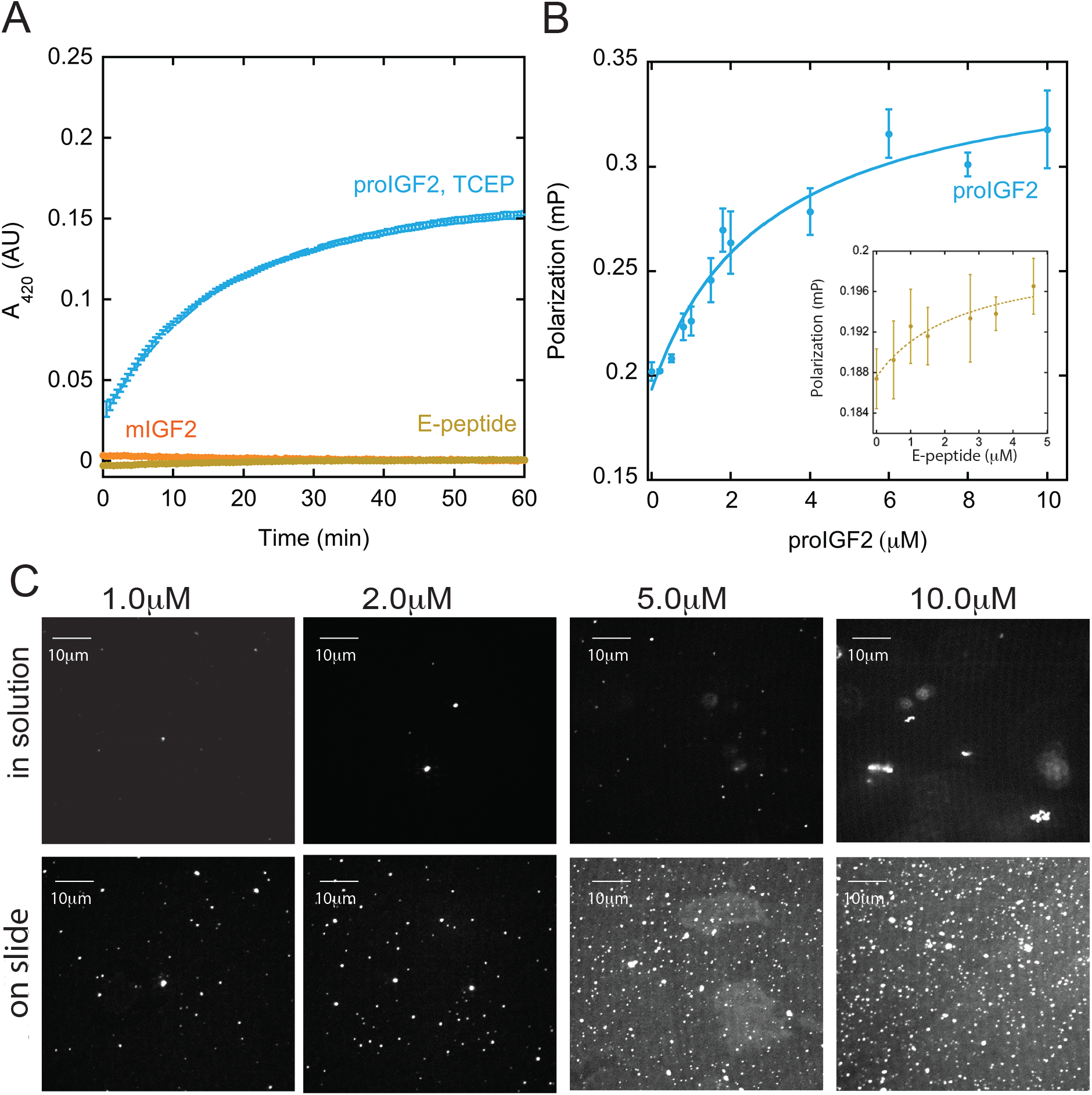
ProIGF2 forms oligomers under reducing conditions. **A)** ProIGF2 produces light scattering under reducing conditions (TCEP), as measured by increased light absorption at 420 nm. Solid line is a single exponential fit (0.057 ± 0.003 min^-1^). **B)** FITC-labeled proIGF2 and E-peptide (inset) were titrated with corresponding constructs under TCEP conditions. Solid line is a single site binding model fit (ProIGF2 *K*_*d,app*_ = 2.98 ± 0.66μM; E-peptide *K*_*d,app*_ =2.5μM). Error bars are the S.E.M of at least three measurements. **C)** Images of Alexa 647-labeled proIGF2 oligomers visualized by confocal microscopy. Buffer conditions: 50mM Tris pH 7.5, 100mM KCl, 1mM ATP, 1mM MgCl_2_, 5mM TCEP, 0.8M urea (proIGF2), 60mM MES pH 6.0, 100mM KCl, 1 mM MgCl2, 1 mM ADP, 0.5 mg/mL BSA, 1 mM DTT (E-peptide);

### ProIGF2 forms oligomers

Under reducing conditions unfolded proIGF2 scatters light, whereas neither the mIGF2 nor the E-peptide region alone scatters light (Figure 4A and Supplemental Figure 5A). Thus, it is the combination of the mature-region and the intrinsically disordered E-peptide region that drives proIGF2 self-assembly.

**Figure 5.**
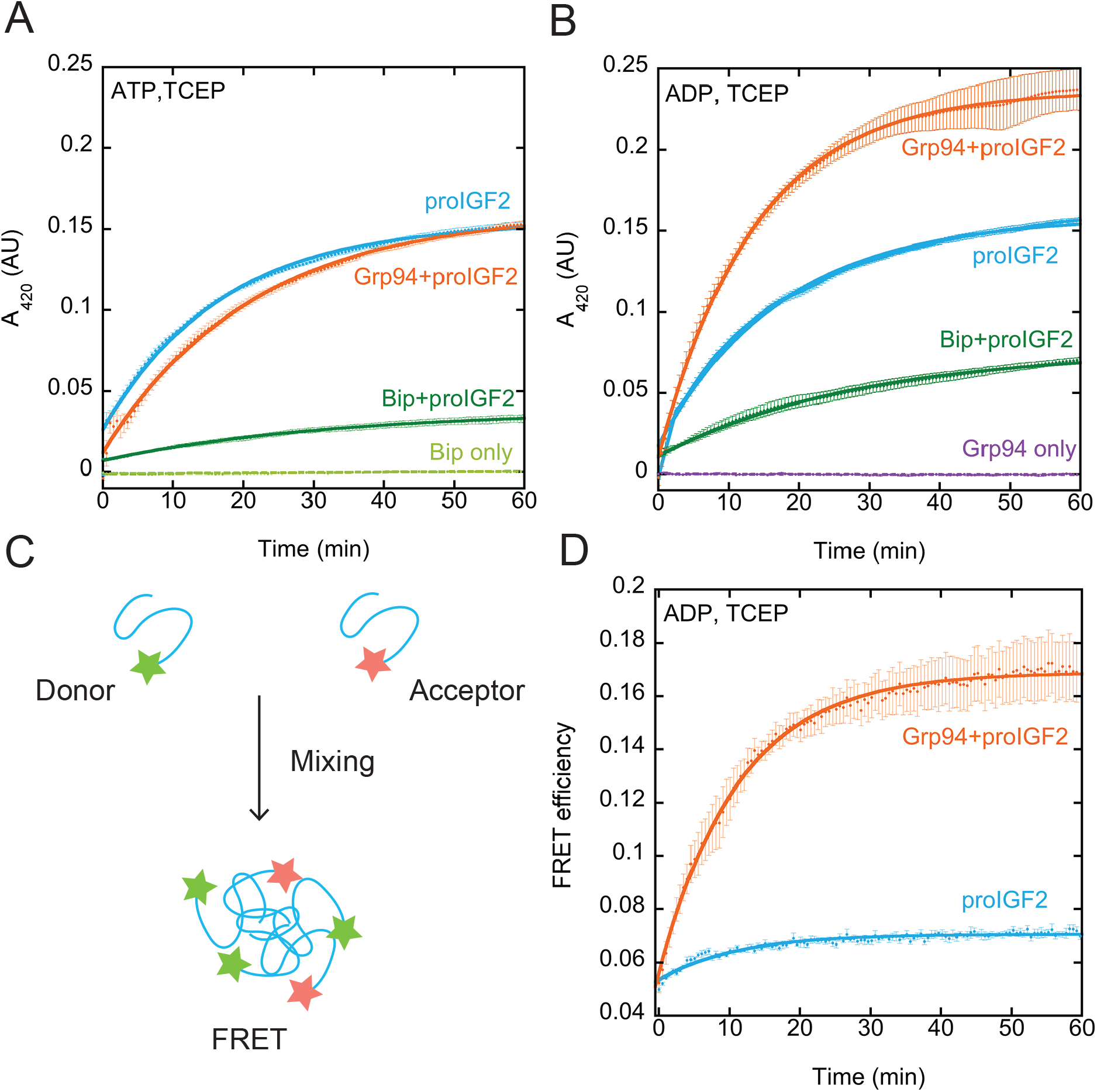
BiP and Grp94 have opposing effect on proIGF2 oliogmerization. **A&B)** ProIGF2 light scattering measurements in the presence of BiP and Grp94 under ATP and ADP conditions. Solid lines are single exponential fits (proIGF2 ATP conditions: 0.057± 0.003 min^-1^; proIGF2 and Grp94 ATP conditions: 0.045± 0.001 min^-1^; proIGF2 ADP conditions: 0.052± 0.001 min^-1^; proIGF2 and Grp94 ADP conditions: 0.074±0.008 min^-1^). **C)** Design of proIGF2 oligomerization FRET measurement. **D)** ProIGF2 FRET in the absense and preasence of Grp94 under ADP conditions. Solid line is a single exponential fit (proIGF2 and Grp94: 0.10± 0.01, proIGF2: 0.12± 0.04 min^-1^). Error bars are the S.E.M of at least three measurements. Buffer conditions: 50mM Tris pH 7.5, 100mM KCl, 1mM ATP /1mM ADP, 1mM MgCl_2_, 5mM TCEP, 0.8M urea.

The light-scattering from proIGF2 oligomerization does not have a lag-phase (Figure 4A), indicating that the self-association does not follow fibril-like kinetics in which rapid growth only occurs after a nucleus is formed. Similarly, proIGF2 self-association as measured by fluorescence-depolarization also does not show evidence of strong cooperativity (Figure 4B), also similar to the behavior of condensates^43^. Dynamic light scattering (DLS) was used to determine the average hydrodynamic radius (R_H_) of proIGF2 oligomers. At early time points with 2μM proIGF2, oligomers have a R_H_ of 230nm and this size increases to 747nm after one hour (Table 1).

**Table 1.**
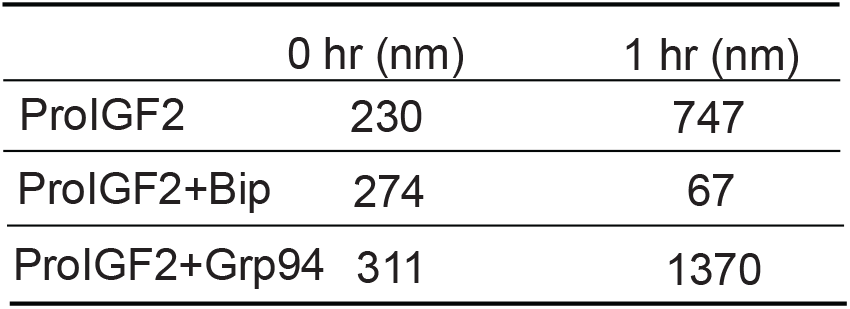
ProIGF2 average particle size (R_H_), measured by DLS under ADP conditions. Measurements were taken directly after oligomer formation (0 hr) and after 1 hour of incubation.

|  | 0 hr (nm) | 1 hr (nm) |
| --- | --- | --- |
| ProIGF2 | 230 | 747 |
| ProIGF2+Bip | 274 | 67 |
| ProIGF2+Grp94 | 311 | 1370 |

Given the critical role of the E-peptide in the formation proIGF2 oligomers, we sought to further characterize this region of proIGF2. Consistent with the sequence disorder prediction in Figure 1A, the E-peptide on its own has minimal secondary structure as measured by circular dichroism (Supplemental Figure 5B). The E-peptide by itself can undergo self-association as measured by fluorescence anisotropy (Figure 4B, inset), which suggests that the oligomerization is driven by weak interactions spread across a largely unstructured E-peptide.

Fluorescently-labeled proIGF2 oligomers can be visualized by confocal fluorescence microscopy (Figure 4C), both in solution and on the microscope slide. These oligomers are predominantly spherical, although irregular shapes are observed at high proIGF2 concentration (see 10μM condition in Figure 4C). Because proIGF2 oligomers can be observed on the slide surface, we tested whether TIRF microscopy could be used to examine the oligomers on more detailed level. The TIRF measurements were performed by forming proIGF2 oligomers at 2.5μM and then diluting proIGF2 to 25nM to achieve sparse coverage on the microscope slide. Under these conditions, proIGF2 oligomers of different sizes are visually apparent (Supplemental Figure 5C) and can be further illustrated by observing variable numbers of photobleaching events on individual proIGF2 oligomers (Supplemental Figure 5D). Quantification of proIGF2 oligomer fluorescence intensity suggests that proIGF2 oligomers have an approximately exponential size distribution rather than forming one specific size (Supplemental Figure 5E).

### BiP and Grp94 have opposing effects on proIGF2 oligomerization

BiP and Grp94 influence proIGF2 oligomerization in very different ways. Under both ADP and ATP conditions BiP suppresses proIGF2 oligomerization (Figure 5A), whereas under ADP conditions Grp94 enhances proIGF2 oligomerization (Figure 5B). Interestingly, Grp94 enhances the magnitude of proIGF2 light-scattering but not the overall kinetics. Neither BiP nor Grp94 produce light scattering on their own (Figure 5A&B). Similar to the optical light scattering, DLS shows Grp94 increasing the R_H_ of ProIGF2 oligomers whereas BiP reduces their size (Table 1). These measurements show BiP decreasing the size of proIGF2 oligomers over time, suggesting that the chaperone can actively disassemble these structures.

The increased ProIGF2 light scattering from Grp94 (Figure 5B) could be explained either by Grp94 enabling proIGF2 oligomers to increase their monomer copy number or alternatively by multiple Grp94 chaperones binding to proIGF2 oligomers and thus increasing the combined particle size. We used FRET to distinguish between these possibilities. We labeled proIGF2 with either a donor or acceptor fluorophore (see Methods). By mixing these separately-labeled samples, proIGF2 oligomerization can be monitored by increasing FRET without any signal interference from Grp94 (Figure 5C). Grp94 enhances proIGF2 oligomerization FRET (Figure 5D) under ADP conditions indicating that Grp94 enables proIGF2 oligomers to increase their monomer copy number. Similar to the light-scattering measurements, we find that Grp94 enhances the extent of proIGF2 oligomerization with a minimal influence on the overall kinetics (Figure 5D).

Interestingly, the above FRET assay demonstrates that proIGF2 slowly transitions from dynamic oligomers to irreversible aggregates. The change in proIG2 oligomer dynamics can be observed by introducing Grp94 to proIGF2 oligomers at different times (Figure 6A). We find that proIGF2 oligomerization FRET can be enhanced by Grp94 and the enhancement of proIGF2 oligomerization decreases over time (Figure 6B). We conclude that dynamic oligomers of proIGF2 slowly transform to irreversible rigid aggregates, and that Grp94 can only act on the dynamic oligomers.

**Figure 6.**
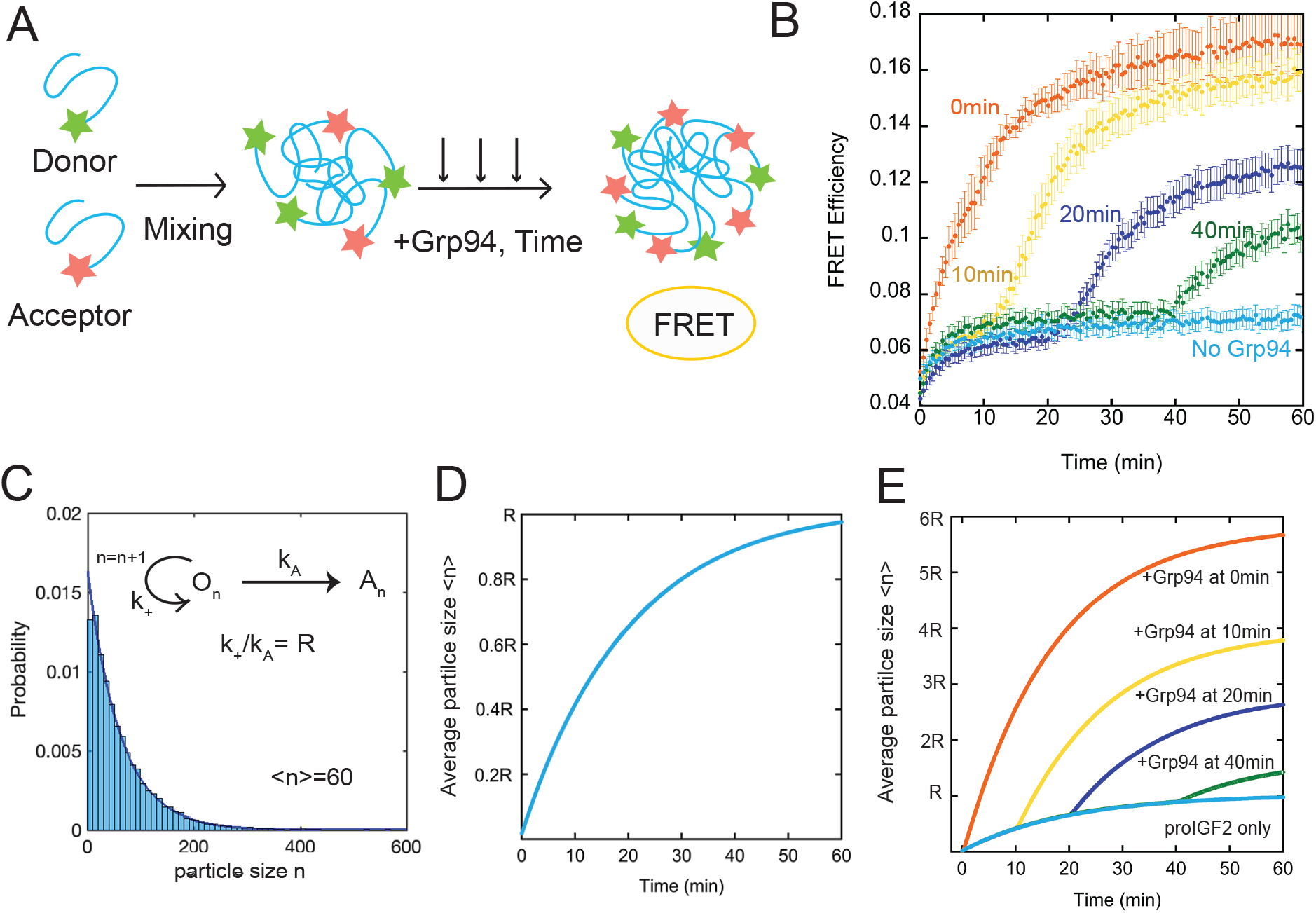
Grp94 targets proIGF2 dynamic oligomers. **A)** Design of proIGF2 oligomerization FRET measurement with Grp94 introduced to proIGF2 oligomers at variable times. **B)** ProIGF2 FRET in the absense and presence of Grp94 under ADP conditions. Error bars are the S.E.M of at least three measurements. Buffer conditions are the same as Figure 5D. **C)** ProIGF2 aggregation and oligomerization model and numerical simulation. Monomers are added to oligomers with a rate of *k*_*+*_ while oligomers transition to aggregates with a rate of *k*_*A*_. Numerical simulation of proIGF2 particle size distribution with *k*_*A*_=0.05min^-1^, *k*_*+*_=3min^-1^ after one hour of oligomer growth. Solid line is a single exponential fit. **D)** Numerical simulation of proIGF2 particle growth over time with *k*_*A*_=0.05min^-1^and *k*_+_=3min^-1^. **E)** Numerical simulations of proIGF2 oligomerization with Grp94 introduced at various times to match the FRET measurements in Figure 6A. The *k*_*+*_ is increased 6-fold by Grp94, therefore the maximium average oligomer size <n> is 6R.

Collectively, the above measurements are consistent with Grp94 targeting proIGF2 oligomers and enhancing their dynamics. Figure 6C presents a minimal quantitative model for proIGF2 oligomerization and aggregation. Monomers of proIGF2 are added with a rate of *k*_*+*_ and oligomers of any size can transition to an irreversibly aggregated state with a rate of *k*_*A*_. The final average size of proIGF2 particles is given by *k*_*+*_/*k*_*A*_ (we denote this ratio as R). This model yields proIGF2 oligomer and aggregate sizes that are exponentially distributed, consistent with the size distribution observed by TIRF microscopy (Supplemental Figure 5E). For example, Figure 6C shows the distribution of proIGF2 particle sizes in which the aggregation rate is set to 0.05 min^-1^ (similar to the aggregation kinetics measured in Figure 4) and the rate of monomer addition is arbitrarily set to 3.0 min^-1^. The resulting particle sizes are exponentially distributed (sold line, Figure 6C), with average size (<n>=60) that is in-line with the expected value (*k*_*+*_/*k*_*A*_=60). This model of oligomer growth does not yield a lag-phase (Figure 6D), consistent with the light-scattering and FRET measurements in Figure 5.

According to the model above, Grp94 can enhance proIGF2 oligomerization either by slowing the aggregation transition (decreasing *k*_*A*_) or by accelerating oligomerization dynamics (increasing *k*_*+*_). Because the light-scattering and FRET both show Grp94 enhancing the magnitude of oligomer formation but not the overall kinetics this implies that Grp94 has a minimal influence on *k*_*A*_ but rather accelerates proIGF2 oligomerization dynamics (Grp94 increases *k*_*+*_). Figure 6D shows an example numerical simulation of the influence of Grp94 on proIGF2 oligomerization in which Grp94 enhances proIGF2 oligomerization dynamics six-fold. In these numerical simulations Grp94 is introduced at various times during the course of proIGF2 oligomer formation, thus matching the design of the FRET experiments in Figure 6B. The numerical simulations and experimental data are in qualitative agreement (compare Figure 6B and Figure 6E). We conclude that the simple model in Figure 6C provides an adequate explanation of the influence of Grp94 on proIGF2 oligomerization.

## Discussion

Here we discover that the BiP and Grp94 chaperones influence both the folding and the oligomerization of proIGF2 (Figure 7). One motivation for studying proIGF2 was to test the idea that Hsp90-family chaperones are specialized for the late stages of protein folding. Although prevalent in the Hsp90 literature^15, 17, 44^ testing this concept has led to diverging views (*e.g.* separate studies of p53 have concluded that client is folded, molten globule, and fully unfolded when bound to Hsp90 ^20-22^.)

**Figure 7.**
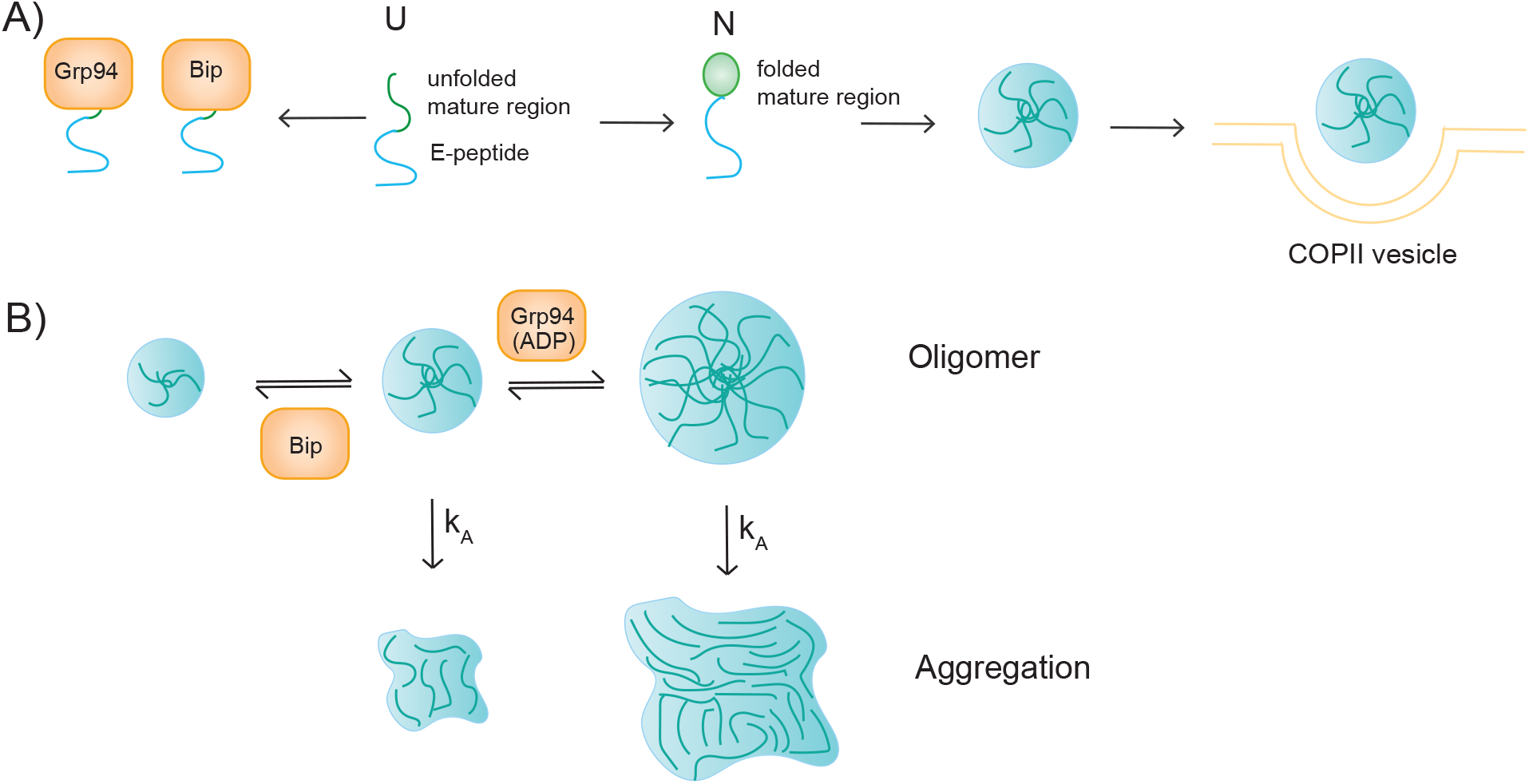
Schematic illustration of Grp94 and BiP chaperone influence on proIGF2. **A)** Both BiP and Grp94 bind the unfolded proIGF2, while the E-peptide region promotes proIGF2 oligomerization facilitating the packaging of proIGF2 into vesicles for transportation to Golgi. **B)** Grp94 in the open conformation (ADP) binds to proIGF2 oligomers and increases their size. BiP decreases the size of proIGF2 oligomers.

An empowering feature of IGF2 is that its disulfide-bonded folding intermediates can be trapped and separated by RP-HPLC similar to classic studies that uncovered the detailed folding pathways of BPTI^29-30^. If Grp94 indeed stabilizes specific folding states of IGF2, this experimental system would be able to detect which state is stabilized and quantify the extent of stabilization. However, we find that both BiP and Grp94 have only a modest influence on the folding properties of proIGF2, primarily by stabilizing the fully unfolded state (Figures 2&3, Supplemental Figure 3). Thus, for the proIGF2 client we do not see evidence that Grp94 is specialized for the later stages of client folding.

In studying the folding properties of proIGF2 and mature IGF2, we discovered that the E-peptide region of proIGF2 promotes oligomerization (Figure 4). The role of the intrinsically disordered E-peptide region in promoting oligomerization is in-line with observations from the field of protein condensates^41, 45-48^. However, while *in-vitro* protein condensates are typically in the range of 10μm in size, proIGF2 oligomers are much smaller (R_H_ ∼200nm).

Many peptide hormones, including insulin, oligomerize to promote efficient sorting in the secretory pathway^49-52^. ProIGF2 oligomerization has a plausible biological role in concentrating the pro-hormone into clusters that can be effectively packaged for transport to the Golgi via COPII vesicles (Figure 7A). Because the E-peptide is cleaved off in the Golgi^34^, the oligomerization property would be relevant to proIGF2 sorting at the ER-to-Golgi transition rather than for proIGF2 folding in the ER. Indeed, we find that the E-peptide has a minimal consequence on the folding of the mature region (Figure 2 & Supplemental Figure 1).

While BiP and Grp94 have a similar influence on proIGF2 folding, these chaperones have opposing effects on proIGF2 oligomerization (Figure 7B). BiP suppresses oligomerization while Grp94 enhances oligomerization (Figure 5). Our findings with Grp94 and proIGF2 are consistent with reports that Grp94 enhances the self-association of the secreted protein myocilin^53^. We find that Grp94 enhances proIGF2 oligomerization by increasing the assembly rate (*k*_*+*_ in Figure 6) rather than from slowing the transition into irreversible aggregates. Similar to Grp94, cytosolic human Hsp90β promotes the oligomerization of the intrinsically disordered protein tau via exposing its aggregation-prone repeat domain^54^. One proposed yet unverified function of chaperones on protein condensates is a role in increasing condensate dynamics as a mechanism to reduce the aging and aggregation process^45^. This idea is in line with our observation that ProIGF2 undergoes a slow transition from dynamic oligomers to irreversible aggregates and that Grp94 can only act on the dynamic oligomers (Figure 6B). More work is needed to explore the physiological consequences of the Grp94 influence on ProIGF2 oligomerization.

Grp94 enhances proIGF2 oligomerization when Grp94 adopts the open state (ADP conditions) but not the closed state (ATP conditions, Figure 4A&B). In a previous study, Ostrovsky and co-workers discovered that ATP binding and hydrolysis are essential for Grp94 to facilitate the cellular secretion of mIGF2^26^. The influence of Grp94 on proIGF2 oligomerization may contribute to the essential ATP-dependent chaperoning function. Future work is needed to determine why the open state of Grp94 exhibits preferential binding proIGF2 oligomers. The Grp94 open state, as with the open state of all Hsp90 chaperones, is conformationally heterogeneous^55-57^. One possibility is that this conformational heterogeneity may help Grp94 interact with the heterogeneous environment of a proIGF2 oligomer.

## Acknowledgements

The authors would like to thank helpful discussion and feedback from Larry Friedman and Street lab members. Research for this project was supported by NIH R01 GM115356 (T.O.S). Authors have no interest of conflict.

## Material and Methods

### Protein expression and purification

Mouse proIGF2, mIGF2 and E-peptide were purified from *Escherichia coli* BL21* cells. Cells were grown at 37°C in LB medium and induced at OD_600_=0.6∼0.8 with 0.1mM IPTG at 30°C overnight. ProIGF2 and mIGF2 proteins were purified through inclusion body extraction and ion exchange chromatography. Purified IGF2 constructs were stored in 8M urea, 100mM Tris pH7.5, 25mM KCl, 1mM EDTA, 1mM TCEP. Purified E-peptide was stored in 8M urea, 25 mM Tris, 250 mM KCl, 1 mM DTT. The Histag of E-peptide was later removed via Histrap column and stored in 25mM Tris pH 7.5, 150mM KCl, 50mM imidazole buffer.

The purification of mouse Grp94 without the charged linker (Δ287-328) and BiP is similar to previously described methods^58-59^. Briefly, BiP and Grp94 were expressed in *Escherichia coli* strain BL21* at 37°C and purified via Histrap column (GE Healthcare), by ion exchange on MonoQ column (GE Healthcare), and by gel filtration on a Superdex S-20 column (GE Healthcare). Proteins were flash-frozen in 50mM Tris, pH7.5, 50mM KCl, 5mM MgCl_2_ and 5% glycerol.

### ATPase measurement

BiP and Grp94 ATPase was measured with a temperature-controlled plate reader (BioTeK) with ATP-regenerating system at 37°C as previously described^60^. All experiments were performed with 50mM Tris, pH 7.5, 100mM KCl, 1mM ATP, 1mM MgCl_2_, 5mM TCEP and 0.8M urea with 1.5μM BiP or 1.5μM Grp94 (dimer), and 5μM IGF2 constructs at 37 °C. ATPase activity was determined by NADH consumption rate measured at 340nm.

### Reverse phase HPLC

To measure proIGF2/mIGF2 folding, samples were diluted 1:10 out of denaturant into refolding/reduced buffer (50mM Tris, pH 7.5, 100mM KCl, 6.25mM GSH, 1.25mM GSSG, or 5mM TCEP (reduced), 1mM ATP/ADP, 1mM MgCl_2_) at 37°C. The final concentrations of proIGF2, BiP and Grp94 were all 5μM. At various time points, proIGF2 refolding reactions were quenched by 500mM H_3_PO_4_ and 5% acetonitrile. The quenched samples were spun down at 15,000g for 3min and the supernatant was injected to C4-300 reverse-phase column (ACE, 100 × 4.6mm, 5μm). Different proIGF2 folding states were separated with 50mM H_3_PO_4_ in H_2_O (Buffer A) with a linear acetonitrile gradient with 50mM H_3_PO_4_ (Buffer B) and a flow rate of 1ml/min. ProIGF2 elution profiles were detected by absorbance at 220nm. To determine the populations of folding states for the three-state analysis (Figure 2D & 1F) the absorbance at 220nm was integrated for the U, I, and N regions and normalized to the total integrated area. The fitting of the three-state model is described in the Appendix.

### Light scattering

IGF2 constructs were diluted to 5μM into a reducing buffer (50mM Tris, pH 7.5, 100mM KCl, 1mM MgCl_2_, 5mM TCEP, 0.8M urea, 1mM ATP/ADP, 25°C) with or without a stochiometric concentration of BiP and Grp94. Light scattering signals were monitored by light absorption at 420nm.

### Dynamic light scattering

ProIGF2 hydrophobic radius (R_h_) was measured with ALV/DLS/SLS-5000 Light Scattering System at room temperature. ProIGF2 samples were diluted to 2μM into a reducing buffer (50mM Tris, pH 7.5, 100mM KCl, 1mM MgCl_2_, 5mM TCEP, 0.8M urea, 1mM ADP, 25°C) with or without 5μM of BiP or Grp94. Samples were spun down at 20000g and the supernatant was monitored at a 90° angle by laser light scattering at 633nm. The translational diffusion coefficient was obtained by analyzing the autocorrelation fluctuation and R_h_ is calculated from the diffusion coefficient. The samples containing chaperones and proIGF2 are polydisperse because R_H_ values of BiP and Grp94 are below 10nm (Bip: 3.0nm, Grp94 3.2nm). Thus, the mass weight distribution (M_w_) of particles with R_H_>10 was excluded to minimize the interference of chaperones.

### Fluorescence polarization measurements

ProIGF2 was labeled with a 2-fold excess of fluorescein isothiocyanate (FITC) for 1hr under buffer conditions of 100mM Tris pH 7.5, 25mM KCl, 1mM EDTA, 8M urea, at room temperature. The labeling reaction was quenched with 5mM 2-mercaptoethanol, and the free dye was removed from labeled protein by ion-exchange chromatography. Protein was stored in 8M urea, 100 mM Tris pH 7.5, 500 mM KCl, 1 mM EDTA, 1 mM TCEP. Fluorescence polarization measurements were performed with 100nM labeled proIGF2 on a Fluoromax-4-spectrofluorometer (Horiba Scientific) with an excitation wavelength of 485nM and emission wavelength of 520nM with 3nm slit width. Labeled proIGF2 was preincubated with proIGF2 in a matching 8M urea buffer before measurements, and then diluted ten-fold into reducing buffer (100mM Tris pH 7.5, 100mM KCl, 1mM ATP, 1mM MgCl_2_, 1mM TCEP, 0.8M urea) to measure the fluorescence polarization at 37°C.

E-peptide was labeled with FITC and fluorescence polarization measurements were performed with 50nM labeled E-peptide titrated by unlabled E-peptide in a reducing buffer (60mM MES pH 6.0, 100mM KCl, 1 mM MgCl2, 1 mM ADP, 0.5 mg/mL BSA, 1 mM DTT). Fluoromax-4-spectrofluorometer (Horiba Scientific) setting is the same as proIGF2 measurements described above.

### Förster resonance energy transfer

ProIGF2 was labeled with a stochiometric concentration of AlexaFluor 555 C2 maleimide or AlexaFluor 647 C2 maleimide (Invitrogen) for 4hr at room temperature. Free dye was removed via an SP column. The labeling rate for both fluorophores was determined to be 20% by spectrophotometric measurements. ProIGF2 was stored in reduced denaturant buffer (100mM Tris pH 7.5, 25mM KCl, 1mM EDTA, 1mM TCEP, 8M urea). FRET measurements were initiated by mixing 250nM donor labeled proIGF2, 250nM acceptor labeled proIGF2 and 2μM wt proIGF2. The mixture was diluted into 25mM Tris pH 7.5, 100mM KCl, 1mM ADP, 1mM MgCl_2_, 5mM TCEP, 0.8M urea buffer. The FRET kinetics were monitored by exciting donor fluorophore at 532nm, and detecting donor and acceptor at 565nm and 670nm respectively, with slit width of 4nm and a time interval of 30s. The FRET efficiency was calculated as *E*_FRET_= *I*_*Acceptor*_*/(I*_*Donor*_*+I*_*Acceptor*_), where *I*_*donor*_ and *I*_*acceptor*_ are the donor and acceptor emission intensities observed at donor and acceptor detection wavelengths.

### Confocal microscopy

Alexa Fluoro-647 labeled proIGF2 was diluted to various concentrations in 50mM Tris pH 7.5, 100mM KCl, 1mM ATP, 1mM MgCl_2_, 5mM TCEP buffer. All proIGF2 oligomer images were obtained on an Observer Z1 microscope (Carl Zeiss) equipped with a CSU-X1 spinning-disk confocal head (Yoka-gawa) and a QuantEM 512SC EM charge-coupled device (CCD) camera (Photometrics) with a 100× (NA 1.45) oil-immersion-objective at room temperature. Confocal Z-stacks were collected every 15s for 5min, with manual focus adjustment. Fluorescence microscope image analysis was performed using ImageJ.

### Total Internal Reflection Fluorescence Microscopy (TIRF)

Alexa Fluoro-555 labeled proIGF2 was diluted from 2.5μM to 25nM into 50mM Tris buffer (pH 7.5, 50mM Tris 50mM KCl, 600μM MgCl_2_, 0.5mg/ml BSA, 600μM ADP and equilibrated at room temperature. Glass slides, coverslips and sample preparation followed previously descriptions^61-62^. Single molecule TIRF imaging of proIGF2 molecule was performed on a custom microscope^63^, using 532nm excitation wavelength and a 2.4s sampling interval with 600μW laser power. Data analysis and spot identification was performed on custom software built in Matlab, available online (https://github.com/gelles-brandeis/CoSMoS_Analysis).

### Numerical simulation

Numerical simulations in Figure 6 were performed in Python using Gillespie sampling. The *k*_*A*_ rate constant (0.05 min^-1^) was determined from the light scattering kinetics in Figure 4. The *k*_*+*_ rate 3 min^-1^ (without Grp94) and 18 min^-1^ with Grp94, were selected arbitrarily. Both oligomers and irreversible aggregations were both included while calculating the average size of proIGF2 particles <n>, which representing the average number of proIGF2 monomer copies in one particle. The average particle size distribution at 1 hour is normalized and fitted to exponential distribution. The average particle size increases with time and are well fitted to exponential rising model, with selected *k*_*A*_ rate (0.05 min^-1^).

### Circular Dichroism

E-peptide was diluted 1:10 out of denaturant at 25°C. The final concentration of E-peptide in the experiment was 8.1 μM in 0.8M urea, 2.5 mM Tris, 25mM KCl, and 0.1mM DTT. Measurements were performed on a Jasco J-810 Circular Dichroism System.

---

**Supplementary Figure 1.**
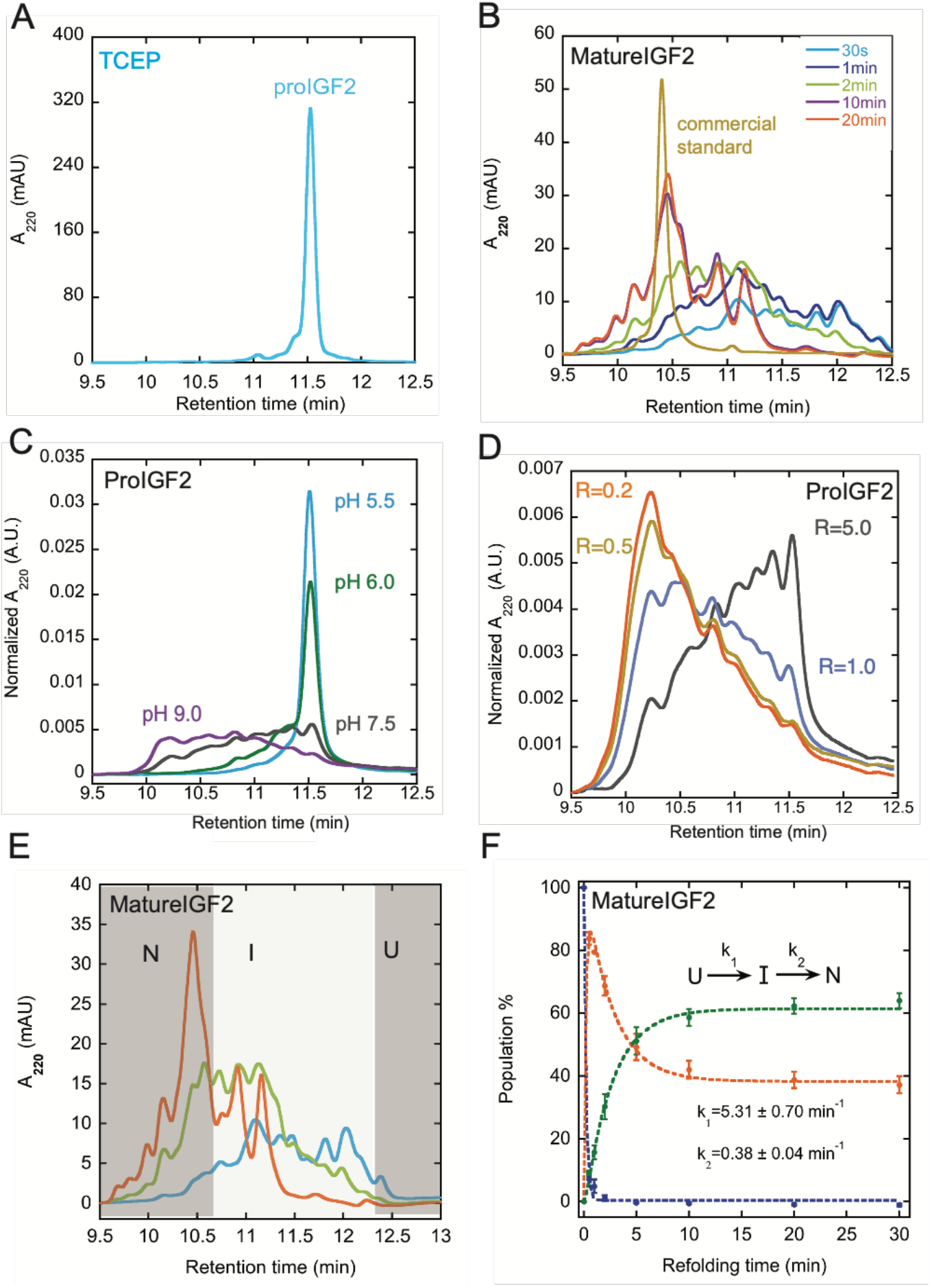
**A)** RP-HPLC analysis of reduced unfolded proIGF2. The fully reduced peak elutes at 11.5min, whereas the oxidized populations elute at earlier times (compare with Figure 2B). **B)** RP-HPLC analysis of mIGF2. A commercial standard of mIGF2 is shown in gold. Refolding times shown in legend. **C&D)** ProIGF2 refolding is pH-dependent and redox potential dependent. The pH-dependent refolding samples and redox-dependent samples were collected at 0.5min. The pH-dependent assay was measured under R=5.0; and the redox-dependent assay was measured under pH 7.5. **E&F)** Mature IGF2 folding is categorized into three states (U, I, and N). Populations of mIGF2 were fitted with a three-state model (see Methods). The populations of U, I and N are color-coded in blue, orange, and green respectively. Error bars are the S.E.M of at least three measurements. Reducing buffer conditions are 50mM Tris pH 7.5, 100mM KCl, 1mM ATP, 1mM MgCl_2_, 5mM TCEP, 0.8M Urea, 37 °C. Refolding buffer conditions are the same as Figure 2. For pH-control assays, buffer reagents from pH 5.5 to 6.5 are MES, and from pH 7.5 to 9.0 are Tris. For redox-control assays, Redox potential R= [GSSG]/[GSH] and GSSG and GSH ratio are adjusted according to experimental conditions.

**Supplementary Figure 2.**
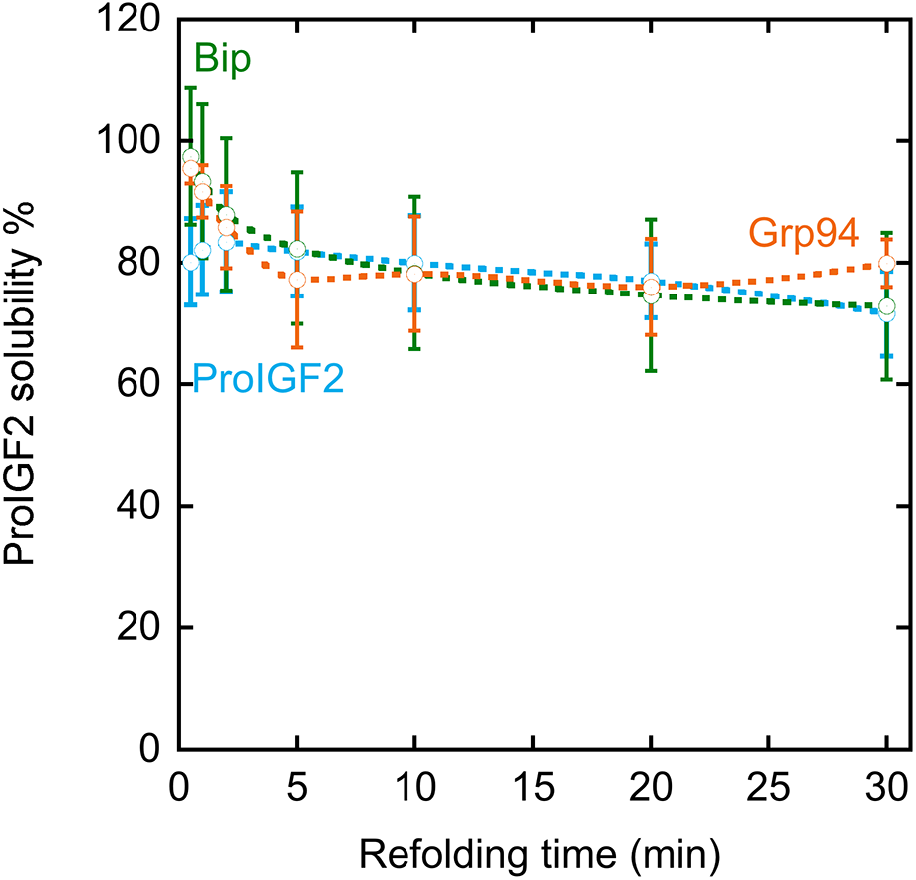
The solubility proIGF2 was calculated by integrating proIGF2 RP-HPLC chromatograms and normalizing these values to the yield of proIGF2 at 0min refolding time. Error bars are the S.E.M of at least three measurements. Refolding buffer is the same as Figure 2.

**Supplementary Figure 3.**
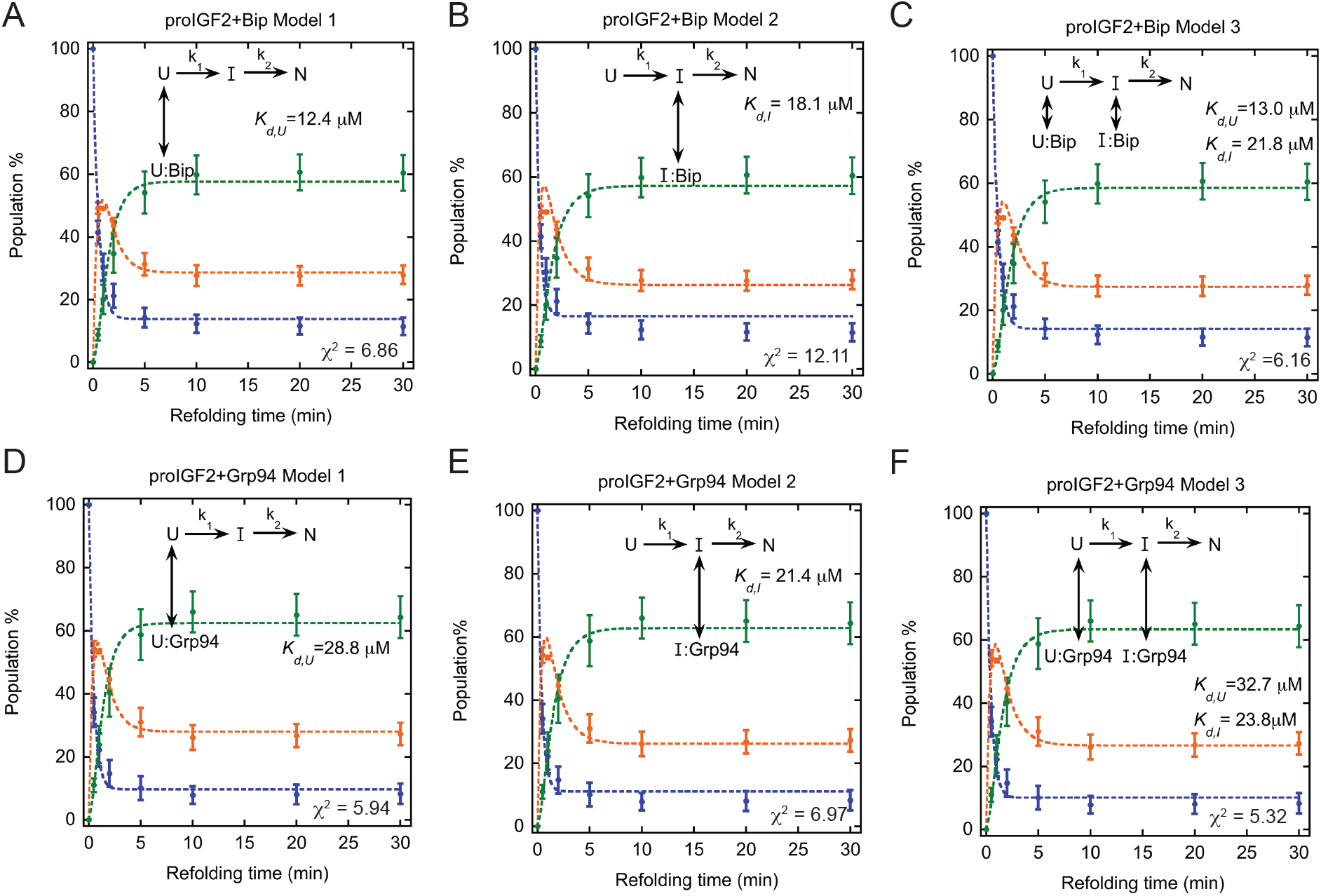
ProIGF2 folding in the presence of BiP (**A-C**) and Grp94 (**D-F**) are fitted to three models. The fitting values of *k*_*1*_ and *k*_*2*_ were obtained from proIGF2 folding in Figure 2D. The binding affinity between BiP/Grp94 to proIGF2 U state and I state were determined according to equations 17&18 in Appendix 1. The fitting results (dashed lines) are compared with experimental data (dots). Unfolded, intermediate and native states are represented in blue, orange and green respectively. More detail information about the model fitting is given in Appendix 1.

**Supplementary Figure 4.**
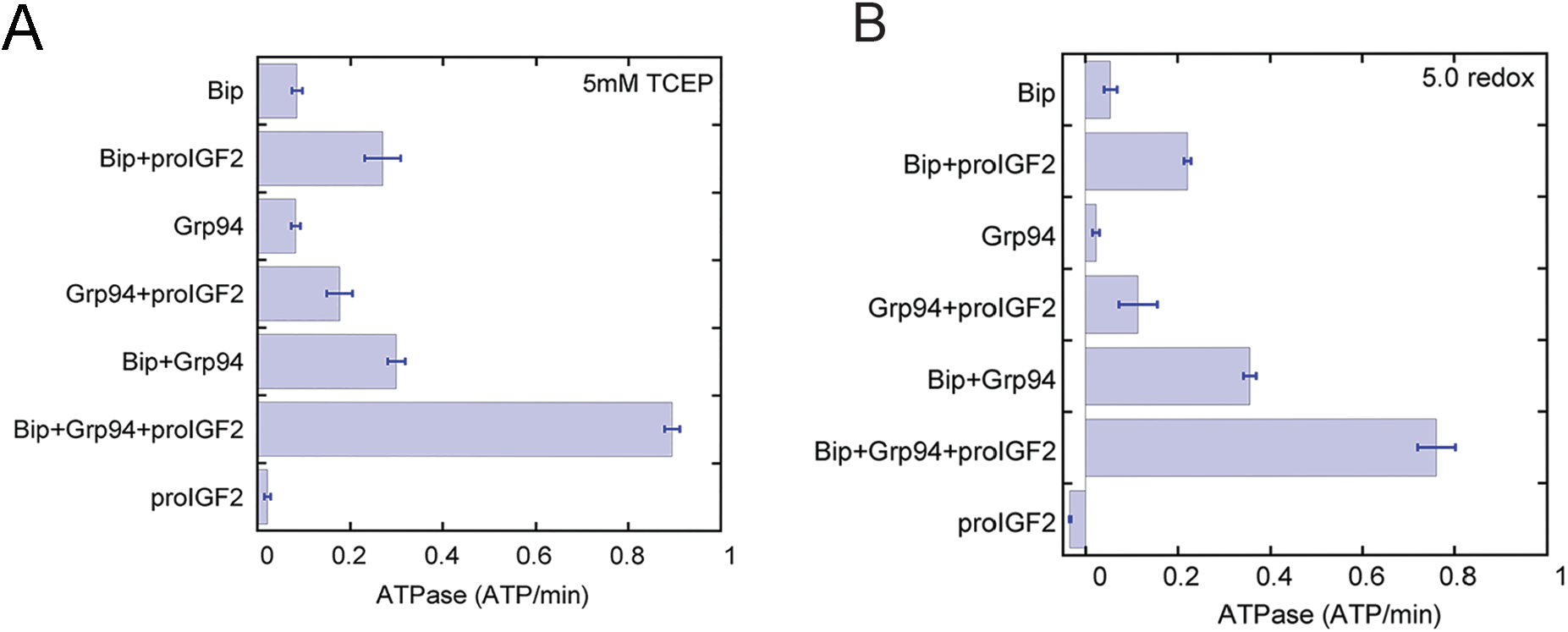
ProIGF2 increases BiP and Grp94 ATPase under reducing conditions **(A)** and refolding conditions **(B)**. Error bars are the S.E.M of at least three measurements. Refolding buffer conditions are the same as Figure 2. Reducing buffer conditions are the same as Supplemental Figure 1A.

**Supplementary Figure 5.**
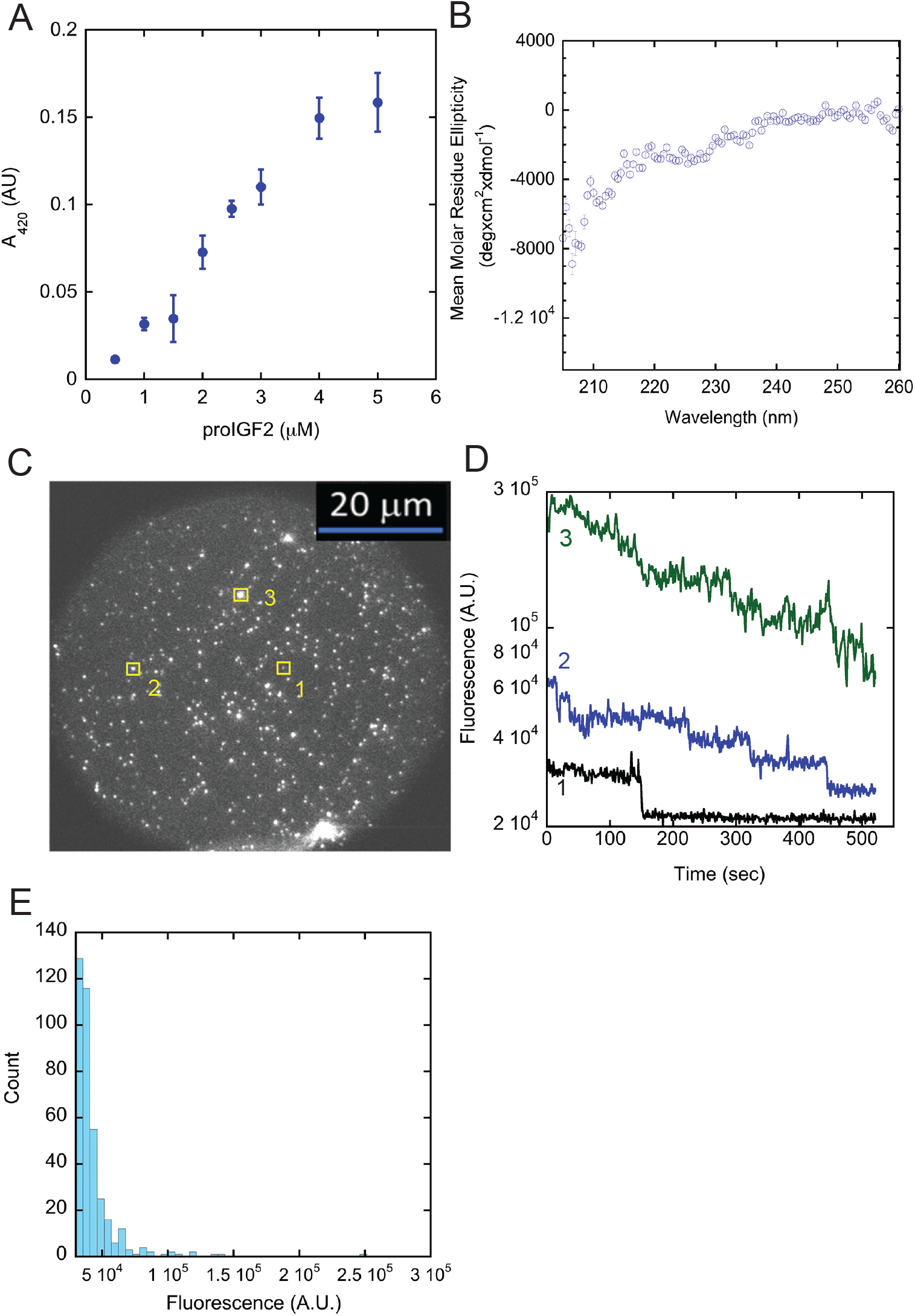
**A)** Light scattering measurements for different concentrations of ProIGF2 measured at 1hr. **B)** Circular dichroism measurement of the proIGF2 E-peptide. The E-peptide has minimal secondary structure, as indicated by the absence of a well-defined minima in the wavelength range of 220 nm. **C)** TIRF microscopy image of fluorescently labeled proIGF2. Three proIGF2 oligomers with distinct sizes are numbered and illustrated in yellow boxes. **D)** Example fluorescence intensity changes for proIGF2 oligomers. Different sizes of proIGF2 oligomers present different photobleaching process. **E)** All spots from the slide in panel C were identified computationally (see Methods) and their integrated fluorescence was calculated, producing the resulting histogram.

