## Appendix for "The ER chaperones BiP and Grp94 regulate the formation of insulin-like growth factor 2 (IGF2) oligomers"

**Appendix: ProIGF2 kinetic model derivations and fitting analysis**

Three-State Model:

We use a three-state kinetic model:

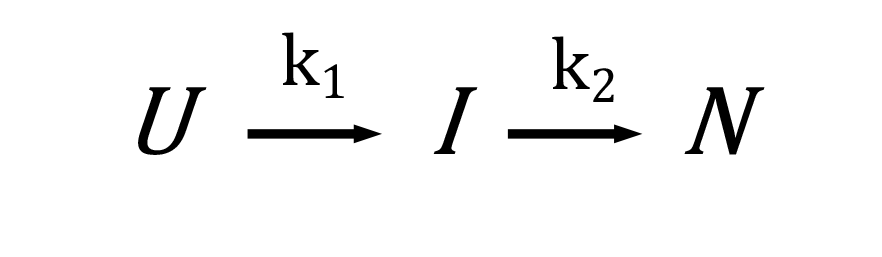

The fraction of unfolded (U), intermediate (I), and native (N) states are given by:

$U=A_{1}\times e^{-k_{1}t} + B_{1}(Eq.1)$

$I=100-(A_{1}\times e^{-k_{1}t} + B_{1})-\{A_{2}\times\{1+\frac{1}{(k_{1}-k_{2})}\times{[k_{2}e}^{(-k_{1}t)}-{k_{1}e}^{(-k_{2}t)}]\} \}(Eq.2)$

$N=A_{2}\times\{1+\frac{1}{(k_{1}-k_{2})}\times{[k_{2}e}^{(-k_{1}t)}-{k_{1}e}^{(-k_{2}t)}]\} (Eq.3)$

Where an additional baseline parameter (*B_1_*) is included to account for residual unfolded population present at long times. Similarly, the *A_2_* parameter accounts for the incomplete conversion of all proIGF2 to the N state.

Chaperone Binding Model #1

This model assumes chaperone (C) binding solely to the unfolded state:

We assume that the binding and unbinding of chaperone is fast compared to the U🡪I transition, thus, this model simplifies to:

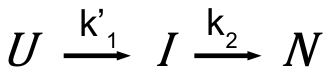

Where

$$k_{1}^{'}=k_{1}\times\frac{K_{d,u}}{(K_{d,u}+[chaperone])} (Eq.4)$$

Chaperone Binding Model #2

This model assumes chaperone binding solely to the intermediate state:

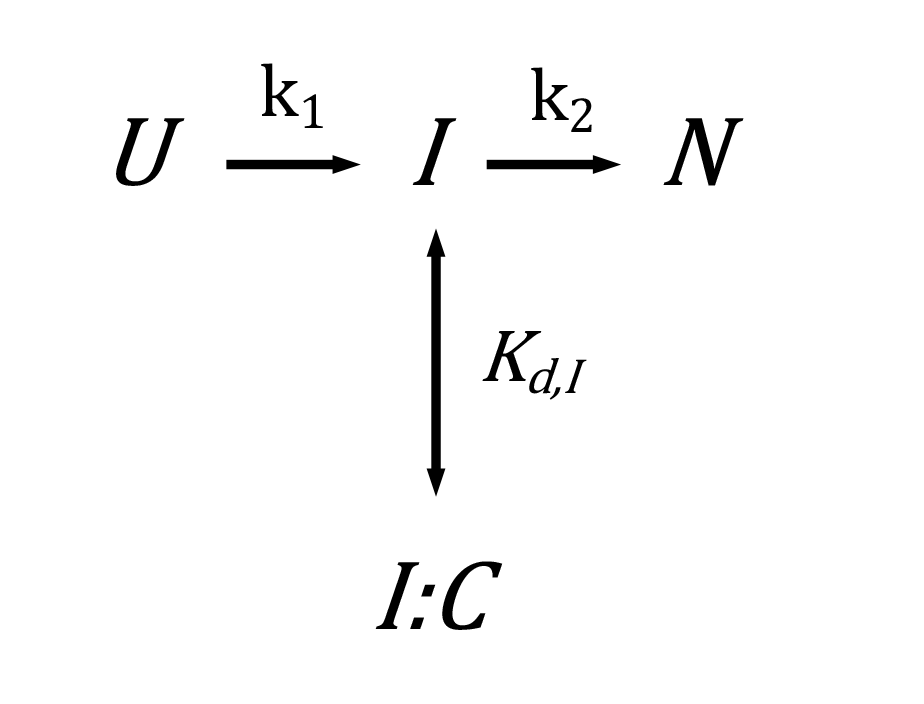

We again assume that the binding and unbinding of chaperone is fast compared to the I🡪N transition, thus, this model simplifies to:

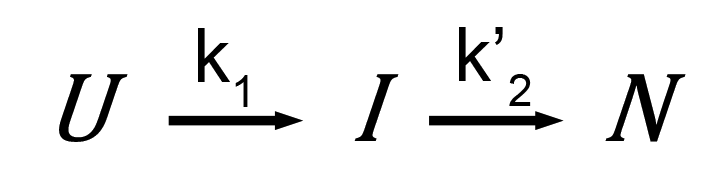

Where

$$k_{2}^{'}=k_{2}\times\frac{K_{d,I}}{(K_{d,I}+[chaperone])} (Eq.5)$$

Chaperone Binding Model #3

This model assumes chaperone binding to the both the unfolded and intermediate states:

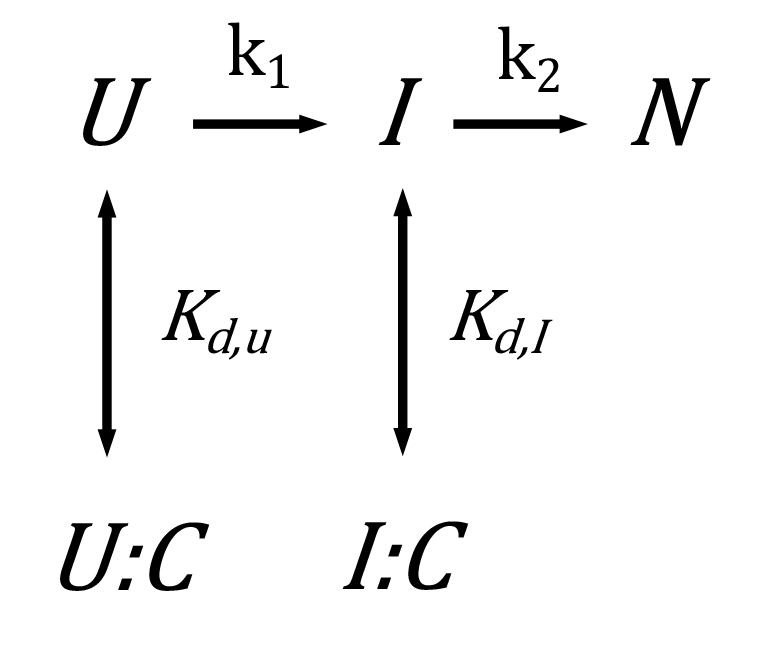

Which simplifies to

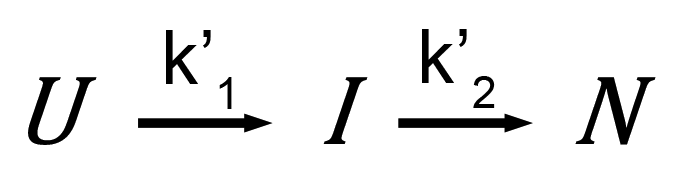

Where $k_{1}^{'}$ and $k_{2}^{'}$ are given by equations Eq.4 and Eq.5.
